# What sets the mutation rate of a cell type in an animal species?

**DOI:** 10.64898/2025.12.19.695482

**Authors:** Marc de Manuel, Molly Przeworski, Natanael Spisak, Anastasia Stolyarova

## Abstract

Germline mutation rates per generation are strikingly similar across animals, despite vast differences in life histories. Analogously, in at least one somatic cell type, mutation rates at the end of lifespan are comparable across mammals. These observations point to a key role for natural selection in shaping mutation rates. In this essay, we summarize the patterns identified to date and outline existing theories for how selection pressures might shape mutation rates in animal germline and soma. We argue that an understanding of what sets the mutation rate of a given cell type in a species requires better integration of genetics and development with population processes of selection and genetic drift.

## MAIN TEXT

Germline mutations are the substrate of evolution. They arise from accidental changes to the genome in a cell lineage leading to egg or sperm and, when transmitted, are present in every cell of the offspring. Their net effect is deleterious (Kimura 1967; Lynch 2010): while most *de novo* mutations are without fitness effects and a tiny fraction are advantageous to the organism, a larger number are instead harmful, introducing perturbations that lead to miscarriages, cause developmental disorders, and contribute to heritable diseases. By contrast, in humans and other species with dedicated germlines, somatic mutations are restricted to a subset of tissues or cells; while most are inconsequential, a subset drives cancer and contributes to the aging process (Alexandrov and Stratton 2014; Vijg and Dong 2020).

Mutations are not only inputs into disease and evolution, however, they are also outputs. Mutations in both germline and soma arise from the interplay between myriad sources of damage, DNA replication cycles, and repair and damage tolerance pathways, involving hundreds of genes (Seplyarskiy and Sunyaev 2021). Human mutation rates are heritable (controlling for parental ages; (Kaplanis et al. 2022; Lawson et al. 2025) and the mutation spectrum evolves over time (e.g., in humans; (Harris 2015; Gao et al. 2023). Viewed through this lens, mutation rates in germline and soma are coupled quantitative traits, subject to genetic drift and natural selection. Understanding the selection pressures that set the mutation rate of a species has been of interest for close to 90 years (Sturtevant 1937; Kimura 1967; Lynch 2010; Peto 2016) and is central to the study of aging as well as of evolution.

For germline mutations, this effort has been motivated by a few “stylized facts”. First, comparisons across prokaryotes, unicellular, and multi-cellular organisms indicate that taxa with larger effective population sizes have lower mutation rates per base pair (bp) per generation (Lynch 2010). Second, mammalian genomes of short-lived species accumulate more mutations per unit time than do those of longer-lived species (Kohne 1970; Wu and Li 1985; Wilson Sayres et al. 2011). Third, in humans and other amniotes, fathers transmit more *de novo* mutations than mothers, reflective of an elevated mutation rate in spermatogenesis relative to oogenesis (Haldane 1947; Crow 2000; de Manuel et al. 2022; Bergeron et al. 2023). These broad strokes have stood the test of time, but can now also be filled in with numerous, direct estimates of germline mutation rates from pedigree studies, as well as mutation rates for diverse tissues and somatic cell types in humans as well as other species.

Here, we summarize the picture that is emerging, notably the striking similarity in germline mutation rates per generation seen across animals, the constancy in mutation rates at the end of lifespan in intestinal crypts, and a seeming deviation from this trend in blood stem cells. We discuss alternative theories about the selection pressures that might give rise to these patterns and how they might be extended to consider differences among cell types.

### Some observations

❖ We collated estimates of germline mutation rates per generation (i.e., at the mean age of reproduction) obtained from whole genome resequencing of pedigrees, requiring a minimum of five trios for the estimate to be relatively precise (Box 1). Although our focus is on mammals, we include additional comparative data from animals to help place the patterns in a larger context. The picture that emerges is remarkable: despite the huge diversity of environments, metabolic rates, gametogenic processes, and three orders of magnitude variation in generation times, germline mutation rates per generation per bp all fall within one order of magnitude (10^-9^ to 10^-8^) (Fig 1). In fact, the same order of magnitude mutation rate per generation per bp is seen even in distantly related multi-cellular organisms, such as mushrooms and plants (Wang and Obbard 2023).
❖ Given that the germline mutation rates per generation are close to constant, it follows that the rate per year is inversely proportional to generation time; we illustrate the pattern in mammals (Fig 2A-B, Fig 3) but it holds for other taxa (including invertebrates) as well (Thomas et al. 2010; Lewin and Eyre-Walker 2025). The observation that long-lived species have a lower mutation rate per unit time is the well known “generation time effect” from evolutionary biology, which has been the object of study since the 1970s (Laird et al. 1969; Kohne 1970; Wu and Li 1985; Britten 1986; Ohta 1992; Martin and Palumbi 1993; Wilson Sayres et al. 2011).
❖ The constancy across species in the per generation mutation rate is all the more striking when one considers that it aggregates contributions from both sexes, different developmental stages, and distinct mutagenic processes. Notably, germline mutations arise early in embryogenesis, in a cell lineage that gives rise to the germline, and during gametogenesis. The relative contributions of these stages differ substantially between short and long-lived species (de Manuel et al. 2022), yet the total mutation rate per bp per generation does not. Similarly, germline mutation rates are an average of maternal and paternal rates. Although the contribution of paternal germline varies slightly among species, within mammals, there is little or no correlation with the total rate (Fig 2C). Moreover, the total mutation rate comprises distinct mutagenic processes, which differ in their proportions across species (Gelova et al. 2022; Cagan et al. 2022; Ramos-Almodóvar et al. 2025).
❖ Compared to somatic cell types, the human germline mutation rate is unusually low, even relative to post-mitotic or quiescent cells: for instance, the male germline accumulates point mutations roughly ten times more slowly than neurons (Abascal et al. 2021). But mutation rates also vary among somatic cell types: for instance, there are an additional 9 mutations per cell per year in bile ducts versus 56 mutations in colonic crypts (Moore et al. 2021).
❖ In the one somatic tissue in which there is comparative *in vivo* mutation data for mammals, colonic epithelium, the same scaling is seen across species as for the germline (Cagan et al. 2022): the mutation rate per year decreases with intrinsic lifespan (which is highly correlated with generation time; (de Magalhães et al. 2007)) (Fig 2D, Fig 3). The same scaling does not appear to hold for blood stem cells, however (Fig 3). Comparing humans and mice, it does not seem to fit fibroblasts or neurons either (Milholland et al. 2017; Hazen et al. 2016).

**Figure 1.**
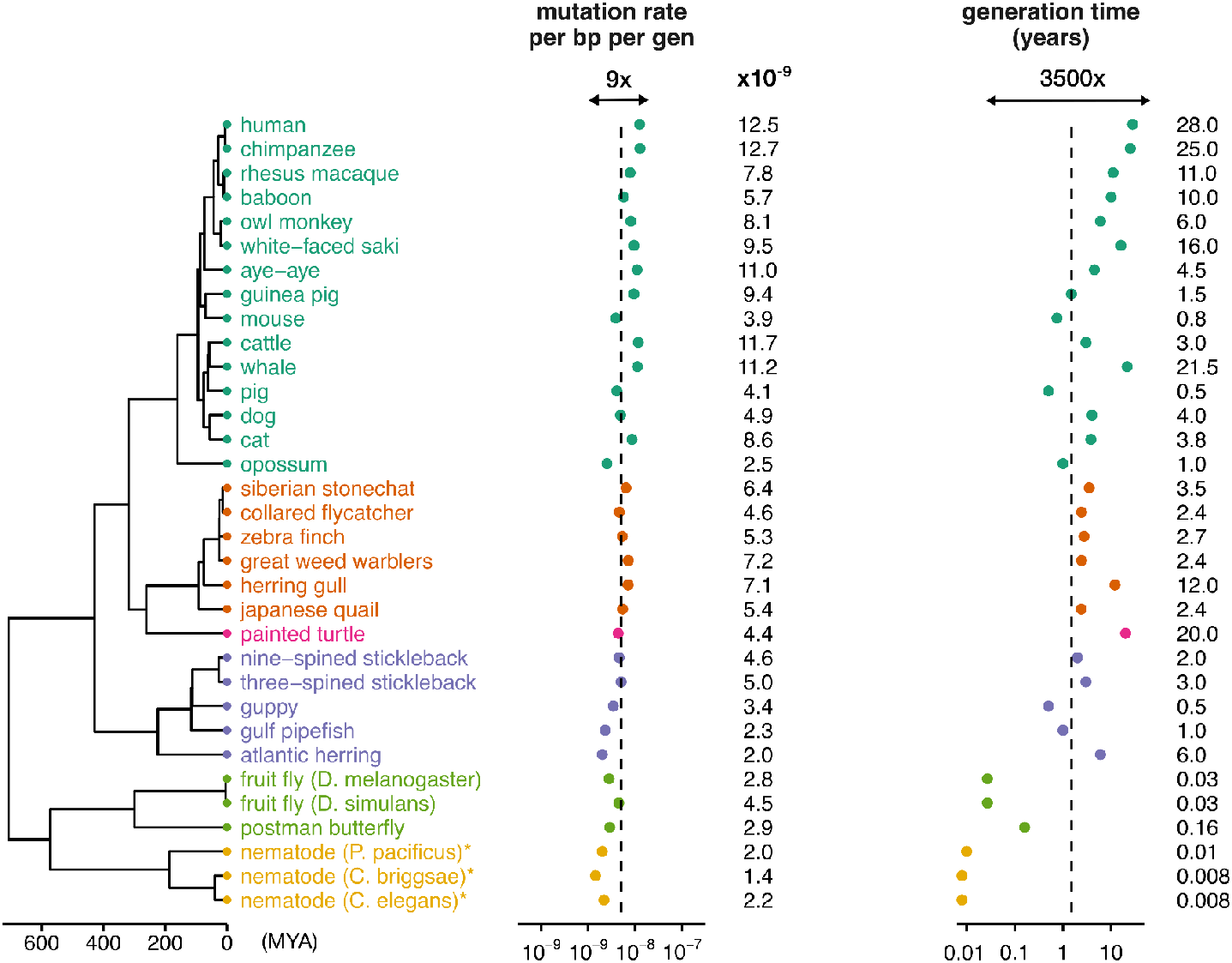
Variation in germline mutation rates per generation across animals. Included are estimates based on a substantial fraction of the genome (in practice, >¼) and at least five trios. All estimates are based on pedigree sequencing, with the exception of mutation-accumulation experiments in worm species (marked with asterisks). Generation time estimates are taken either from the original pedigree studies or, when not provided, from the AnAge database of animal life-history traits (de Magalhães et al. 2024). For species with multiple independent estimates, we report the average mutation rate weighted by the number of trios in each study. All data sources are listed in Supplementary Table 1.

**Figure 2.**
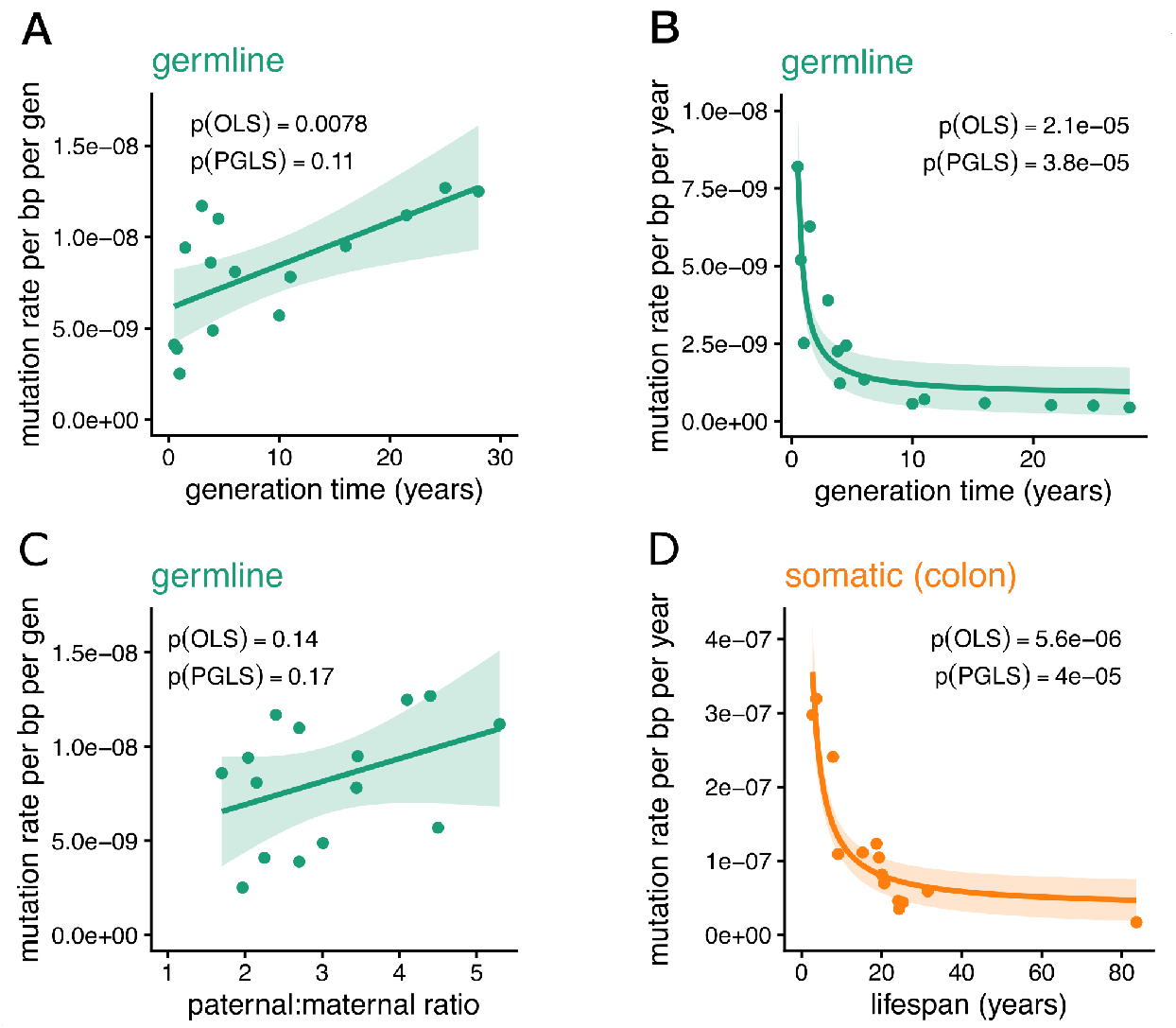
Relationships of mutation rates to generation times and paternal bias in mutation in mammals. Based on 15 mammalian species, using the same data in Figure 1. Reported are p-values for an OLS and PGLS (phylogenetic generalized least squares) regressions (Freckleton et al. 2002). Trend lines are from the OLS regression. **(A)** The germline mutation rate per bp generation shows little or no increase with generation time. **(B)**. The germline mutation rate per bp per year scales inversely with the generation time. **(C)** No relationship is detectable between paternal bias in mutation and the germline mutation rate per bp per generation. **(D)** The mutation rate per bp per year in colonic crypts scales inversely with the lifespan. Intrinsic lifespan estimates are based on (Cagan et al. 2022).

**Figure 3.**
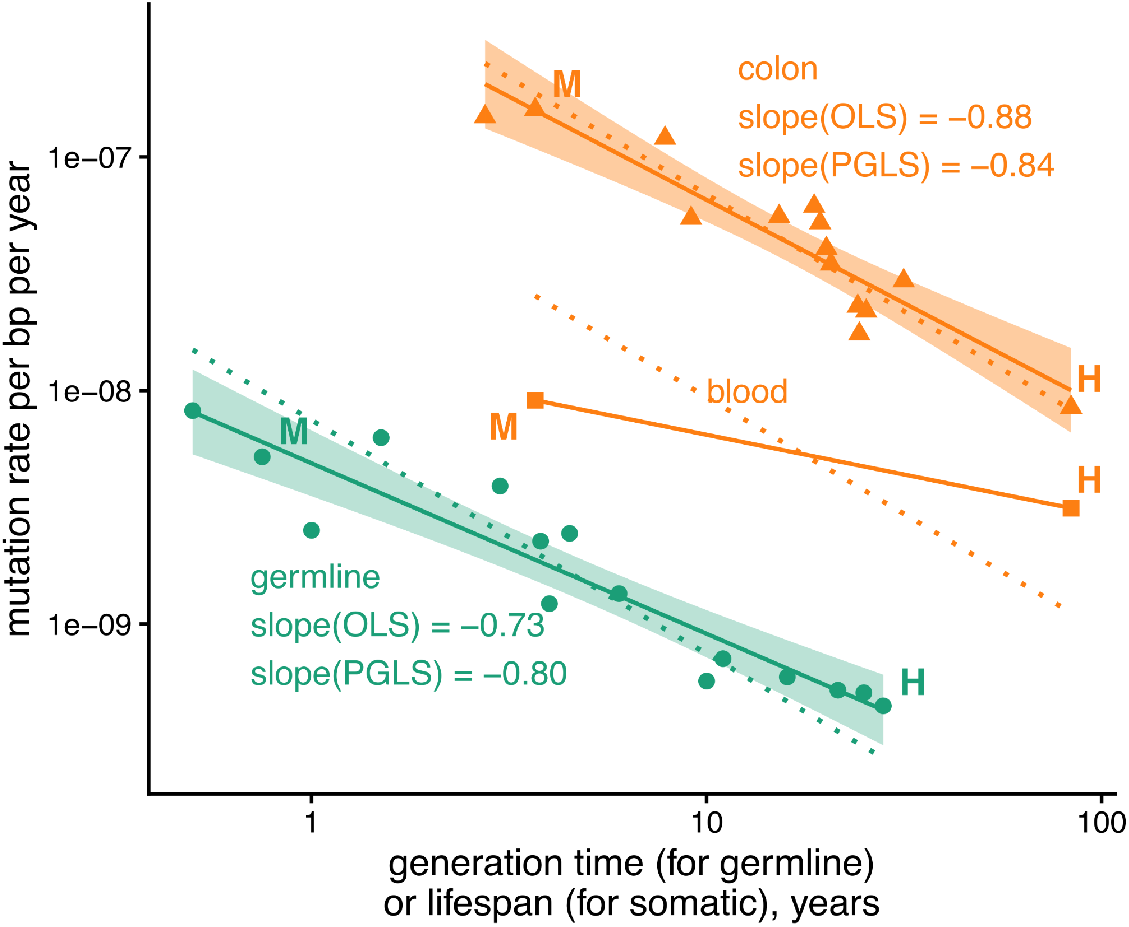
Relationship of the mutation rate per year to the generation time or lifespan on a log-log scale. In orange triangles are data from colonic crypts (Cagan et al. 2022) and in orange squares those for blood stem cells (Mitchell et al. 2022; Kapadia et al. 2025) (for human H and mouse M), while in blue dots are germline mutation rates (for males and females combined, as in Fig 1). Solid lines show the fitted regression slopes obtained from OLS; dotted lines indicate a slope of -1, which corresponds to mutation rates scaling perfectly with the inverse of generation time or lifespan.

In summary, the mutation rate per generation appears to fall between 10^-8^ and 10^-9^ per bp across animals, despite changes in all its constituent components (Fig 1; Box 1). A small range of mutation rates is also seen across mammals in colonic crypts at the end of lifespan (Fig 3) (Cagan et al. 2022). Such constancy is not expected from genetic drift alone, given the vast phylogenetic distances represented, nor is it likely to arise as a simple byproduct of life history or developmental constraints, since these differ dramatically among species. Instead, the similarity points to strong selection pressures on the mutation rate per bp per generation (or lifespan). As expected if the net effect of a mutation is deleterious, there appears to be directional selection to decrease the mutation rate as much as possible. The specific value of the rate in a species may be set by limits to the efficacy of selection (see Box 2); alternatively, driving the rate down even further may inflict other costs, e.g., on developmental timing, such that optimum is under (indirect) stabilizing selection (Kondrashov 1995).

Regardless, the relatively invariant mutation rate per generation/lifespan (Fig 1) and the slower mutation rate per unit time in long-lived mammals (Fig 2B, 2D) are two facets of the same picture. What is not clear is whether the picture emerges from selection on the rate per generation, per year, or both.

### Some theories

#### The view from the germline

In evolutionary biology, models for the evolution of germline mutation consider selection on modifier alleles that increase the number of mutations inherited by offspring every generation (and have no other effects) (Kimura 1967; Lynch 2008). Because on average a germline mutation is deleterious, such modifiers are selected against. Consistent with this expectation, the few mutator alleles mapped to date in mammals are at very low frequency in the population (Kaplanis et al. 2022; Stendahl et al. 2023; Sherwood et al. 2023; Young et al. 2024; Andrianova et al. 2024) and the heritability of human mutation rates is inferred to lay among rare variants (Kaplanis et al. 2022). The efficacy with which selection weeds mutator alleles out depends on their fitness costs and on the extent of genetic drift in the species (see Box 2). Assuming all else is equal, at equilibrium, sexually-reproducing species with higher levels of genetic drift are predicted to have a higher mutation rate per generation (Lynch 2010).

All is not equal, however: notably, species with high levels of genetic drift also tend to have longer generation times (Ohta 1992). If modifier alleles introduce mutations per year (rather than per generation), then they should inflict larger fitness costs in such long-lived species, compared to shorter lived species with more effective selection. Taking this feature into account, modeling suggests that mutation rates per generation may increase only slightly with longer generation times (and greater genetic drift; (Zhu et al. 2025)). This prediction appears to fit the observations within animals (Figs 1 & 2A). However, the fact that it fits mutation rates per bp, rather than per genome, requires further explanation (Box 2).

If strong selection maintains a relatively constant per generation mutation rate, it follows that the rates per year will be inversely proportional to the generation time (Fig 2B), with no need to invoke additional selection pressures on rates per unit time. Observations in colonic crypts (Fig 2D) could then be explained either through pleiotropic effects of the repair and damage tolerance machinery active in the germline or because similar selection pressures shape (some types of) somatic cells.

#### The view from the soma

Somatic mutations are the primary cause of cancers and are thought to contribute to aging by a variety of pathogenic mechanisms (Failla 1958; Szilard 1959; Martincorena and Campbell 2015; Balmain 2020; Lodato and Walsh 2019; Vijg and Dong 2020; Kennedy et al. 2012; Kirkwood 2005). Notably, mutations can increase rates of cell proliferation, leading to clonal expansions that cause signaling problems, compromise tissue function, or lead to cancer (Jaiswal and Ebert 2019; Watson et al. 2020; Alon 2023; Yoshida 2025). Long-standing multistage models of carcinogenesis (Armitage 1985; Stratton et al. 2009) posit that cancers develop once a stem cell has experienced K driver mutations, where K depends on the tissue or cell type, reflecting differences in the physiological environment and the impact of tissue architecture on cancer initiation and promotion. Even in cases where driver mutations are not sufficient causes of cancer (Rozhok and DeGregori 2015) (or even necessary causes; (Balmain 2023)), they greatly increase the risk.

In turn, somatic mutations have long been hypothesized to drive aging (Szilard 1959). A recent theory (Karin et al. 2019; Alon 2023) proposes that the Gompertz law of mortality (i.e., the observation that the mortality rate increases exponentially with age) results from saturation in the removal of damage with age. In this framework, mutations in stem cells are considered a source of damage that gives rise to impaired differentiated cells, ultimately resulting in the accumulation of senescent cells and compromising tissue function.

If selection acts to limit carcinogenesis and the rate of aging, we would expect lower somatic mutation rates per year in longer-lived species. Accordingly, associations have been reported between the efficiency of several DNA repair pathways and life history traits in mammals (Hart and Setlow 1974; Tian et al. 2019; Firsanov et al. 2025) and the yearly somatic mutation rate in colonic crypts decreases with intrinsic lifespan (Cagan et al. 2022) (Fig 2D). Depending on coevolved life strategies among species, selection pressures may depend on intrinsic lifespan or on lifespan in the wild (Alon 2023). In species with high extrinsic mortality, the somatic mutation rate will only be under selection to the limited extent that it leads to early deaths (Cagan et al. 2022). In contrast, in long-lived species with late reproduction and low fecundity or which inhabit an environmentally protected niche (Szekely et al. 2015; Alon 2023), selection on mutation rates may contribute to shaping lifespan.

Existing observations about somatic mutations across mammals are consistent with the notion that selection acts on the fitness effects of somatic mutations. If so, the analogous observations for germline mutations (Fig 1B) could be explained by pleiotropic effects on mechanisms of genome maintenance. However, if selection stems primarily from the effects of somatic mutations, it remains unclear why germline mutation rates are so much lower than those observed in somatic tissues.

#### A possible synthesis

In reality, mutation rates are likely under some degree of selection in both somatic and germline cell types: for instance, the risk of cancer is greatly increased by some modifiers of germline mutation rates (Kaplanis et al. 2022; Robinson et al. 2021). Moreover, not all somatic cell types are created equal: the effect of a mutation to organismal fitness presumably depends on the cell type function and its susceptibility to malignant transformation. Consider a simplified world view, in which there are four types of cells in the body:

A. Long-lived stem cells, which can give rise to cancer or compromise tissue function via mutations (among other sources of damage). In these cells, there is a selection pressure to decrease the mutation rate per unit time, to the extent allowed by the drift barrier.
B. Transient cells that are only present in development (e.g., neural progenitor cells, embryonic stem cells). In this case, it is the fitness costs of mutations that arise during this transient period that matters.
C. Differentiated cells that are rapidly and regularly replaced (e.g., neutrophils, skin fibroblasts), in which direct selection pressures on mutation rates are likely low.
D. Fully differentiated cells that do not get replaced in adulthood (e.g., neurons, cardiomyocytes, beta cells) and thus must persist over much of the lifespan of an individual. In these types, as in (A), selection is presumably acting on the mutation rate per unit time.

In this view, cell types leading to gametes belong to multiple categories but are a special case in which selection is especially strong, because the mutations they carry become constitutive in the offspring.

Somatic cells, meanwhile, occupy different positions along this continuum, depending on their rate of replacement and contribution to organismal fitness. Therefore, we would expect mutation rate to be lowest in germline and highest in short-lived, differentiated cells, with somatic stem cells lying in between.

In this respect, it is intriguing that the patterns observed across mammals (notably the slopes in Fig 3) point to similar selection pressures on the germline and colonic crypts across species. Specifically, the observed relationships between mutation rates per unit time and generation time (or lifespan) are consistent with the fitness cost of mutators being directly proportional to generation time (or lifespan) and the same across mammals (Zhu et al. 2025). A similar scaling does not seem to hold for all cell types, however. In blood stem cells, mice exhibit a lower yearly mutation rate than expected relative to humans (or equivalently, there is an unexpectedly high rate in humans, Fig 3). This observation may be explained by different selection pressures on mutations in the two species, arising from distinct effects of mutations on cancer risk (e.g., because blood cancer is initiated by fewer driver mutations or more readily in mice than humans (de Ruiter et al. 2018)), a larger numbers of sites in mice at which mutations compromise function, or stronger pleiotropic effects of mutation modifiers on other cell types in mouse.

### Possible mechanistic underpinnings

Recent evidence suggests that most point mutations in germline and soma can be explained by two ubiquitous COSMIC mutational signatures: single bp substitution signature SBS1 (which consists in transitions at methylated CpGs, due to deamination and/or replication errors) and an SBS5-like flat signature (potentially caused by collateral errors during translesion synthesis and repair) (Rahbari et al. 2015; Moore et al. 2021; Ramos-Almodóvar et al. 2025; Spisak et al. 2025). Both of these signatures are generally “clock-like” in that they accumulate linearly with age (Alexandrov et al. 2015; Spisak et al. 2024). Other mutational processes are not well captured by COSMIC signatures, such as the specific C>G substitutions enriched in aging oocytes in apes (Jónsson et al. 2017). In several somatic cell types, there is also evidence of other known damage-induced processes, such as mutations due to oxidative damage or UV radiation (Cagan et al. 2022; Moore et al. 2021). Overall, in the vertebrate soma and germline, mutations in long-lived cells, such as types (A) and (D) described above, appear to be predominantly caused by DNA damage and increase linearly over time (Alexandrov et al. 2015; Spisak et al. 2024; Jaiswal and Ebert 2019). We note, however, that a fraction of mutations carried by a cell originate during short-lived cell states, such as embryogenesis (B), or in rapidly replaced differentiated types (C).

In principle, for selection to reduce the mutation rate in a cell type, it could act to increase repair accuracy (e.g., the error rate of a polymerase; (Lynch 2011)), increase repair efficiency (e.g., the speed of repair enzymes), or decrease DNA damage rates (e.g., by increasing melanin production in skin). Because a number of lesion types lead to mutations owing to error-prone genome replication, slowing down cell division may also increase repair efficiency, all else being equal (Seplyarskiy et al. 2018). In addition to preventing the accrual of mutations, selection can target mechanisms to reduce the consequences of mutations, e.g., by promoting the removal of mutated cells (e.g., (Behm et al. 2023)), by evolving multi-layered anti-tumorogenic mechanisms (e.g., (Rangarajan et al. 2004; Sulak et al. 2016)), or by developing tissue architectures that reduce the chance of uncontrolled clonal expansions (e.g., (Derényi and Szöllősi 2017)).

Some strategies for reducing the mutation rate will be shared among cell types, while others may be specific to a cell type or tissue. If a few key genes influence the repair accuracy, changes to such genes will likely be highly pleiotropic, influencing mutation rates across many cell types. This broad impact increases the fitness cost of modifiers that decrease accuracy, making them more likely to overcome the drift barrier (Box 2). In contrast, modifiers of DNA damage rates may be restricted in their effects to specific cell types or tissues, given that DNA damage likely depends on cellular programs and metabolism, as well as on exposures to mutagens that are environment-specific. Thus, depending on the role of the cell, there may be varying levels of selection to prevent the effects of DNA damage, and in some species, greater functional costs to doing so (e.g., increasing melanin could decrease vitamin D synthesis; (Brenner and Hearing 2008)). These considerations suggest that differences in repair accuracy across species may fit the predictions of the drift-barrier hypothesis (Box 2), whereas the extent of DNA damage in any given cell type, and the selection pressures to reduce it, may be more variable among species.

### Outlook

To date, theories about selection pressures on mutation have been developed largely in parallel in different fields, with evolutionary biologists focused on the germline and emphasizing the role of population processes of selection and drift, and cancer biologists and aging specialists highlighting the deleterious consequences of mutations in the soma. Moreover, while existing models often rely on assumptions that mutations arise from replication errors or track cell divisions (e.g., (Lynch 2008; Tomasetti et al. 2017; Alon 2023)), recent evidence suggests that most mutations in germline and soma are triggered by endogenous and exogenous sources of damage and can arise independently of cell divisions (Gao et al. 2019; Seplyarskiy and Sunyaev 2021; Abascal et al. 2021; Spisak et al. 2024, 2025). There is therefore a need for new theory, which integrates these recent findings and knowledge of tissue architecture into models for selection on mutation rates, and contends with the pleiotropic effects that couple aspects of germline and somatic mutagenesis. Ultimately, predicting the mutation rate of a cell type in a given species likely requires a framework that includes the developmental trajectory of the cell, its longevity, and other properties that influence its contribution to organismal fitness. For a specific cell type, the selection pressures on modifier alleles will depend on features like the mutational target size that can drive cancer or lead to incorrect sensing of hormone levels as well as on pleiotropic consequences on other cell types.

As suggested by evolutionary models, the “scaling law” observed in the germline and colonic crypts of mammals, in which mutation rates per year are inversely proportional to generation time (or lifespan) (Fig 3), is expected to arise when mutations inflict a similar fitness cost per year across species. Not all cell types are expected to meet these conditions, however: in some, mutations are likely inconsequential and mutation rates will only be constrained by pleiotropic effects; in others, the fitness consequences of mutations will differ among species, or the main influence will come from modifier alleles that introduce mutations per generation rather than per unit time. Consequently, we predict that as somatic mutation data becomes available for a wider range of species and tissues, deviations from this scaling law will emerge. Conversely, we should learn a great deal about selection pressures shaping the different cell types from such comparative data.

## Acknowledgments

We thank Pierre Vanderhaeghen for helpful discussions and Ziyue Gao, William Milligan and Jonathan Pritchard for helpful comments on the manuscript. This work was supported by R35 GM083098 to MP and by Ramón y Cajal fellowship RYC2022-037185-I to MdM.

## Box 1

**Estimates of mutation rates**

Germline mutation rates are typically estimated by resequencing the genomes of parents and their offspring, usually from blood-derived DNA. After applying filters based on allelic balance, read support, and other quality measures, researchers identify alleles present in roughly half the reads from the offspring but absent from parental genomes (Roach et al. 2010; Keightley et al. 2014; Bergeron et al. 2023). Standard trio sequencing captures constitutive germline mutations and some early embryonic mutations in the child, but tends to filter out early embryonic or gonosomal mutations in the parent (Moorjani, Gao, et al. 2016). Nevertheless, comparisons with single-cell and single-molecule sequencing in humans suggest that trio-based estimates are reliable (Wang et al. 2012; Moore et al. 2021; Spisak et al. 2024).

Several complementary approaches have emerged that enable mutations to be assigned to different stages of the germline. Designs include sequencing of DNA from sperm, which directly samples the paternal germline (Harland et al. 2017; Shoag et al. 2025; Neville et al. 2025) and sequencing blood-derived DNA of twins or three-generation pedigrees, which improves sensitivity to early embryonic mutations (Sasani et al. 2019; Jonsson et al. 2021; Porubsky et al. 2025).

A number of factors should lead estimates of the per-generation mutation rate to be noisier than they are in reality. *De novo* mutations are rare events, so studies based on a small number of trios inevitably have large sampling error. Variation in parental ages, especially paternal age, strongly affects the number of transmitted mutations, adding to the heterogeneity across datasets. In addition, studies differ in their filtering criteria and quality-control thresholds, which can introduce further variability (Bergeron et al. 2022). At the same time, filtering pipelines are often somewhat *ad hoc* and may be tuned, intentionally or not, to produce mutation rates that look reasonable. Such choices can make estimates appear more similar across studies or species than they actually are. Nonetheless, the overall consistency is unlikely to be an artifact, given that phylogenetic substitution-based analyses show a similar pattern (Wilson Sayres et al. 2011; Moorjani, Amorim, et al. 2016; Lewin and Eyre-Walker 2025).

In turn, somatic mutation rates can be estimated by sequencing single cells (recently reviewed in (Shao et al. 2025)), *in vitro* clones (Milholland et al. 2017), or in specific settings, bulk sequencing of primary tissues. For example, (Cagan et al. 2022) sequenced individual clonal crypts, which should represent clonal outgrowths of single stem cells. Because this approach relies on bulk sequencing of the clone, it primarily captures mutations that arose in the founding stem or early progenitor cells rather than in fully differentiated cells.

## Box 2

**The drift-barrier hypothesis and a recent extension**

Since the average effect of a germline mutation is deleterious, on balance an individual would be better off with no new mutations (Kimura 1967). Why then do germline mutations even occur; why does natural selection not drive the mutation rate to zero? Sturtevant (1937), who first posed this question, speculated that perhaps it is simply not possible for an organism to prevent all mutations.

As Lynch (2008) pointed out, however, the answer may instead lie in population genetic processes. In a small, sexually-reproducing population with lots of genetic drift, a mutator allele that leads to a small number of additional mutations per generation may not inflict enough of a fitness cost on the organism for that modifier to be effectively selected against; in other words, natural selection may not overcome the “drift barrier”. As the effective population size (often denoted *N*_e_) increases, however, so will the efficacy of selection against the modifier allele. Thus, all else being equal, at equilibrium, species with larger *N*_e_ will have lower germline mutation rates per generation. Consistent with this expectation, per bp per generation mutation rates vary by two to three orders of magnitude between single cell eukaryotes with huge *N*_e_ and vertebrates, in which *N*_e_ is much smaller (Lynch 2010; Lynch et al. 2023).

As well appreciated, species with different *N*_e_ also vary in a number of other potentially relevant factors. Notably, animal species with smaller *N*_e_ tend to live longer (Ohta 1992): within mammals, consider a gorilla versus a rabbit for instance. Imagine a modifier that introduces mutations, not per generation, but per year (a “clock-like” modifier). Then, as Zhu et al. (2025) point out, even though there is more drift in gorillas than rabbits, the fitness cost of the modifier will be larger in gorillas, because it will cause more mutations by reproduction. Specifically, they assume that the fitness cost of a clock-like modifier is fixed across species and proportional to the generation time. The net effect of these two countervailing effects–greater drift and greater selection–depends on the precise relationship between effective population sizes and generation times. But they could plausibly lead mutation rates per generation to be relatively constant or to increase only slightly with *N*_e_ values (Zhu et al. 2025).

In this modeling framework, the fitness effect of a modifier allele depends on the cumulative cost of the excess deleterious mutations that it generates across the genome. Therefore, a relative constancy of mutation rates *per bp* across species further requires that the number of functionally-important base pairs in the genome be comparable. Although this assumption is unlikely to hold exactly, it may be approximately the case: despite vast differences in the genome sizes across animals, there is roughly 2-fold variation in the number of exonic base pairs (excluding species with recent whole genome duplications) (Ensembl release 115 (Dyer et al. 2025)).

This modeling framework further assumes that the modifier has as its only phenotypic consequence the introduction of additional mutations. In reality, it could occasionally have beneficial pleiotropic consequences that outweigh its mutagenic effects; for example, the modifier could lead to a new cellular program that is advantageous but increases DNA damage rates. More generally, as formulated to date, the drift-barrier hypothesis focuses primarily on the germline and the fitness effects of mutations transmitted to the offspring of carriers (but see (Lynch 2008)). As the authors of these theories note, however, modifiers of germline mutation rates are likely to also affect somatic mutation rates, imperiling the viability of their carriers (Lynch et al. 2023; Beichman et al. 2024). To integrate selection pressures on both germline and soma, we need to better understand the extent to which mutation rates are coupled across cell types.

